# Large-Scale Genome-Wide Meta Analysis of Polycystic Ovary Syndrome Suggests Shared Genetic Architecture for Different Diagnosis Criteria

**DOI:** 10.1101/290502

**Authors:** Felix Day, Tugce Karaderi, Michelle R. Jones, Cindy Meun, Chunyan He, Alex Drong, Peter Kraft, Nan Lin, Hongyan Huang, Linda Broer, Reedik Magi, Richa Saxena, Triin Laisk-Podar, Margrit Urbanek, M. Geoffrey Hayes, Gudmar Thorleifsson, Juan Fernandez-Tajes, Anubha Mahajan, Benjamin H. Mullin, Bronwyn G.A. Stuckey, Timothy D. Spector, Scott G. Wilson, Mark O. Goodarzi, Lea Davis, Barbara Obermeyer-Pietsch, André G. Uitterlinden, Verneri Anttila, Benjamin M Neale, Marjo-Riitta Jarvelin, Bart Fauser, Irina Kowalska, Jenny A. Visser, Marianne Anderson, Ken Ong, Elisabet Stener-Victorin, David Ehrmann, Richard S. Legro, Andres Salumets, Mark I. McCarthy, Laure Morin-Papunen, Unnur Thorsteinsdottir, Kari Stefansson, 23andMe Research Team, Unnur Styrkarsdottir, John Perry, Andrea Dunaif, Joop Laven, Steve Franks, Cecilia M. Lindgren, Corrine K. Welt

**Affiliations:** MRC Epidemiology Unit, University of Cambridge, United Kingdom; The Wellcome Trust Centre for Human Genetics, University of Oxford, Oxford, UK; Department of Biological Sciences, Faculty of Arts and Sciences, Eastern Mediterranean University, Famagusta, Cyprus; Center for Bioinformatics & Functional Genomics, Department of Biomedical Sciences, Cedars-Sinai Medical Center, Los Angeles, CA; Division of Reproductive Endocrinology and Infertility, Department of Obstetrics and Gynaecology, Erasmus University Medical Centre, Rotterdam, The Netherlands; Department of Internal Medicine, University of Kentucky College of Medicine; University of Kentucky Markey Cancer Center; Departments of Epidemiology and Biostatistics, Harvard T.H. Chan School of Public Health; Department of Internal Medicine, Erasmus University Medical Centre, Rotterdam, The Netherlands; Estonian Genome Center, University of Tartu, Tartu, Estonia; Broad Institute of Harvard and MIT and Massachusetts General Hospital, Harvard Medical School, Boston, United States of America; Department of Obstetrics and Gynaecology, University of Tartu, Estonia; Division of Endocrinology, Metabolism, and Molecular Medicine, Department of Medicine, Northwestern University Feinberg School of Medicine, Chicago, Illinois; Center for Genetic Medicine, Northwestern University Feinberg School of Medicine, Chicago, Illinois; Department of Anthropology, Northwestern University, Evanston, Illinois; deCODE genetics/Amgen, Reykjavik, Iceland; Oxford Centre for Diabetes, Endocrinology and Metabolism, University of Oxford, Oxford; Department of Endocrinology & Diabetes, Sir Charles Gairdner Hospital, Nedlands, Western Australia; School of Medicine and Pharmacology, University of Western Australia, Crawley, Western Australia; Keogh Institute for Medical Research, Nedlands. Western Australia; Department of Twin Research & Genetic Epidemiology, King’s College London, London, UK; Division of Endocrinology, Diabetes and Metabolism, Department of Medicine, Cedars-Sinai Medical Center, Los Angeles, USA; Department of Medicine, Division of Genetic Medicine, Vanderbilt University Medical Center, Nashville, Tennessee, United States of America; Vanderbilt Genomics Institute, Vanderbilt University Medical Center, Nashville, TN; Division of Endocrinology and Diabetology, Department of Internal Medicine Medical University of Graz, Austria; Stanley Center for Psychiatric Genetics, Broad Institute of MIT and Harvard, Cambridge, Massachusetts, USA; Massachusetts General Hospital and Harvard Medical School, Boston, USA; Department of Epidemiology and Biostatistics, MRC-PHE Centre for Environment and Health, School of Public Health, Imperial College London, London, United Kingdom; Center for Life Course Health Research, Faculty of Medicine, University of Oulu, 90014 Oulu, Finland; Biocenter Oulu, P.O. Box 5000, Aapistie 5A, FI-90014, University of Oulu, Finland; Unit of Primary Care, Oulu University Hospital, Kajaanintie 50, P.O. Box 20, FI-90220 Oulu 90029 OYS, Finland; Department of Reproductive Medicine and Gynaecology, University Medical Center Utrecht, The Netherlands; Department of Internal Medicine and Metabolic Disorders, Medical University of Białystok, Białystok, Poland; Department of Internal Medicine, Section of Endocrinology, Erasmus University Medical Centre, Rotterdam, The Netherlands; Odense University Hospital, University of Southern Denmark, 5000 Odense C, Denmark; Department of Physiology and Pharmacology, Karolinska Institutet, Stockholm, Sweden; Department of Medicine, Section of Adult and Paediatric Endocrinology, Diabetes, and Metabolism, The University of Chicago, Illinois, United States of America; Department of Obstetrics and Gynecology and Public Health Sciences, Penn State University College of Medicine, Hershey, Pennsylvania, United States of America; Competence Centre on Health Technologies, Tartu, Estonia; Institute of Clinical Medicine, Department of Obstetrics and Gynecology, University of Tartu, Tartu, Estonia; Institute of Bio- and Translational Medicine, University of Tartu, Tartu, Estonia; Department of Obstetrics and Gynecology, University of Helsinki and Helsinki University Hospital, Helsinki; Oxford NIHR Biomedical Research Centre, Churchill Hospital, Oxford, UK; Department of Obstetrics and Gynecology, University of Oulu and Oulu University Hospital, Medical Research Center, PEDEGO Research Unit, Oulu, Finland; Faculty of Medicine, University of Iceland, 101 Reykjavik, Iceland; 23andMe, Inc., Mountain View, CA, 94041; Division of Endocrinology, Diabetes and Bone Disease, Icahn School of Medicine at Mount Sinai, New York, NY 10029; Institute of Reproductive & Developmental Biology, Department of Surgery & Cancer, Imperial College London, London, United Kingdom; Division of Endocrinology, Metabolism and Diabetes, University of Utah, Salt Lake City, UT, 84112; Reproductive Endocrine Unit, Massachusetts General Hospital, Boston, Massachusetts

## Abstract

Polycystic ovary syndrome (PCOS) is a disorder characterized by hyperandrogenism, ovulatory dysfunction and polycystic ovarian morphology. Affected women frequently have metabolic disturbances including insulin resistance and dysregulation of glucose homeostasis. PCOS is diagnosed with two different sets of diagnostic criteria, resulting in a phenotypic spectrum of PCOS cases. The genetic similarities between cases diagnosed with different criteria have been largely unknown. Previous studies in Chinese and European subjects have identified 16 loci associated with risk of PCOS. We report a meta-analysis from 10,074 PCOS cases and 103,164 controls of European ancestry and characterisation of PCOS related traits. We identified 3 novel loci (near *PLGRKT, ZBTB16 and MAPRE1*), and provide replication of 11 previously reported loci. Identified variants were associated with hyperandrogenism, gonadotropin regulation and testosterone levels in affected women. Genetic correlations with obesity, fasting insulin, type 2 diabetes, lipid levels and coronary artery disease indicate shared genetic architecture between metabolic traits and PCOS. Mendelian randomization analyses suggested variants associated with body mass index, fasting insulin, menopause timing, depression and male-pattern balding play a causal role in PCOS. Only one locus differed in its association by diagnostic criteria, otherwise the genetic architecture was similar between PCOS diagnosed by self-report and PCOS diagnosed by NIH or Rotterdam criteria across common variants at 13 loci.

---

Polycystic ovary syndrome (PCOS) is the most common endocrine disorder in reproductive aged women, with a complex pattern of inheritance^1–5^. Two different diagnostic criteria based on expert opinion have been utilized: The National Institutes of Health (NIH) criteria require hyperandrogenism (HA) and ovulatory dysfunction (OD)^6^ while the Rotterdam criteria include the presence of polycystic ovarian morphology (PCOM) and requires at least two of three traits to be present, resulting in four phenotypes (**Supplementary Figure 1**)^6, 7^. PCOS by NIH criteria has a prevalence of ∼7% in reproductive age women worldwide^8^; the use of the broader Rotterdam criteria increases this to 15-20% across different populations^9–11^.

PCOS is commonly associated with insulin resistance, pancreatic beta cell dysfunction, obesity and type 2 diabetes (T2D). These metabolic abnormalities are most pronounced in women with the NIH phenotype^12^. In addition, the odds for moderate or severe depression and anxiety disorders are higher in women with PCOS^13^. However, the mechanisms behind the association between the reproductive, metabolic and psychiatric features of the syndrome remain largely unknown.

Genome-wide association studies (GWAS) in women of Han Chinese and European ancestry have reproducibly identified 16 loci^14–17^. The observed susceptibility loci in PCOS appeared to be shared between NIH criteria and self-reported diagnosis^17^, which is particularly intriguing. Genetic analyses of causality (by Mendelian Randomization analysis) among women of European ancestry with self-reported PCOS suggested that body mass index (BMI), insulin resistance, age at menopause and sex hormone binding globulin contribute to disease pathogenesis^17^.

We performed the largest GWAS meta-analysis of PCOS to date, in 10,074 cases and 103,164 controls of European ancestry diagnosed with PCOS according to the NIH (2,540 cases and 15,020 controls) or Rotterdam criteria (2,669 cases and 17,035 controls), or by self-reported diagnosis (5,184 cases and 82,759 controls) (**Table 1** and **Supplementary Table 1**). We investigated whether there were differences in the genetic architecture across the diagnostic criteria, and whether there were distinctive susceptibility loci associated with the cardinal features of PCOS; HA, OD and PCOM. Further, we explored the genetic architecture with a range of phenotypes related to the biology of PCOS, including male-pattern balding^18–21^.

**Table 1.** Characteristics of PCOS cases and controls from each cohort included in the meta-analysis.

| Cohort | Subject Type | Number | Age (years) | BMI (kg/m<sup>2</sup>) | PCOS Definition | HA<sup>(1)</sup> n(%) | OD n(%) | PCOM n(%) |
| --- | --- | --- | --- | --- | --- | --- | --- | --- |
| Rotterdam | Cases** | 1184 | 28.8 (4.8) | 26.1 (6.3) | NIH (41%) & Rotterdam (100%)<sup>(2)</sup> | 439 (37.0) | 946 (79.8) | 661 (55.8) |
|  | Controls | 5799 | 60.5 (7.9) | 27.6 (4.7) | Population Based Rotterdam Study | NA | NA | NA |
| UK (London/Oxford) | Cases** | 670 | 32.1 (6.8) | 28.2 (7.9) | NIH (33%) & Rotterdam (100%)<sup>(2)</sup> | 455 (67.9) | 537 (80.1) | 383 (57.2) |
|  | Controls | 1379 | 45 (0)<sup>§</sup> | 26.8 (5.5) | 1958 British Birth Cohort | NA | NA | NA |
| EGCUT | Cases** | 157 | 30.7 (8.2) | 26.2 (6.7) | Rotterdam<sup>(2)</sup> | NA | NA | NA |
|  | Controls | 2807 | 31.5 (7.3) | 23.1 (5.5) | Population Based | NA | NA | NA |
| deCODE | Cases** | 658 | 41.3 (8.7) | 30.1 (7.8) | NIH (56%) & Rotterdam (100%)<sup>(2)</sup> | 644 (97.9) | 380 (57.7) | 507 (77.1) |
|  | Controls | 6774 | 49.0 (9.9) | 25.1 (4.9) | Population Based | NA | NA | NA |
| Chicago | Cases* | 984 | 28.6 (5.5) | 35.9 (8.5) | NIH | 984 (100) | 984 (100) | NA |
|  | Controls | 2963 | 46.8 (15.2) | 27.0 (7.4) | Population Based NUGene | NA | NA | NA |
| Boston | Cases* | 485 | 28.4 (6.7) | 30.8 (8.7) | NIH | 485 (100) | 485 (100) | 441 (90.9) |
|  | Controls | 407 | 27.2 (6.5) | 23.8 (4.1) | Screened controls<sup>(3)</sup> | 0 | 0 | 177 (43.4) |
| 23andMe | Cases*** | 5,184 | 45.1 (13.6) | 29.2 (8.2) | Self report (defined by questionnaire) | NA | NA | NA |
|  | Controls | 82,759 | 51.1 (15.7) | 26.1 (6.1) | No PCOS by self report (defined by questionnaire) | NA | NA | NA |

(1) Clinical or Biochemical.
(2) Rotterdam diagnostic criteria include the NIH criteria. All subjects from the indicated cohorts were used in the Rotterdam analysis.
(3) Controls were screened for regular menses and no hyperandrogenism.
PCOS diagnosis was based on NIH criteria (\*), Rotterdam criteria (\*\*), or self report (\*\*\*). Results are reported as mean (SD) or a number (%). Abbreviations: BMI: body mass index, NA: not available, HA: hyperandrogenism, OD: ovulatory dysfunction, PCOM: polycystic ovarian morphology. <sup>§</sup>All subjects are from the British Birth Cohort (born in 1958).

We identified 14 genetic susceptibility loci associated with PCOS, adjusting for age, at the genome-wide significance level (P < 5.0 x 10^-8^) bringing the total number of PCOS associated loci to nineteen (**Table 2**, **Figure 1**). Three of these loci were novel associations (near *PLGRKT, ZBTB16* and *MAPRE1,* respectively; shown in bold in **Table 2**). Six of the 11 reported associations were previously observed in Han Chinese PCOS women^14, 15^. Eight loci have been reported in European PCOS cohorts^16, 17^. Obesity is commonly associated with PCOS and in most of the cohorts cases were heavier than controls (**Table 1**). However, adjusting for both age and BMI did not identify any novel loci; and the 14 loci remained genome-wide significant. Only one SNP near *GATA4/NEIL2* showed significant evidence of heterogeneity across the different diagnostic groups (**Figure 2**; **Table 2**, rs804279, Het P=2.9×10^-4^). For this SNP, the largest effect was seen in NIH cases and the smallest in self-reported cases. Credible set analysis, which prioritises variants in a given locus with regards to being potentially causal, was able to reduce the plausible interval for the causal variant(s) at many loci (**Supplementary Table 2**). Of note, 95% of the signal at the *THADA* locus came from two SNPs. Examination of previously published genome-wide significant loci from Han Chinese PCOS^14, 15^ demonstrated that index variants from the *THADA, FSHR, C9orf3, YAP1* and *RAB5B* loci were significantly associated with PCOS after Bonferroni correction for multiple testing in our European ancestry subjects (**Supplementary Table 3**).

**Figure 1.**
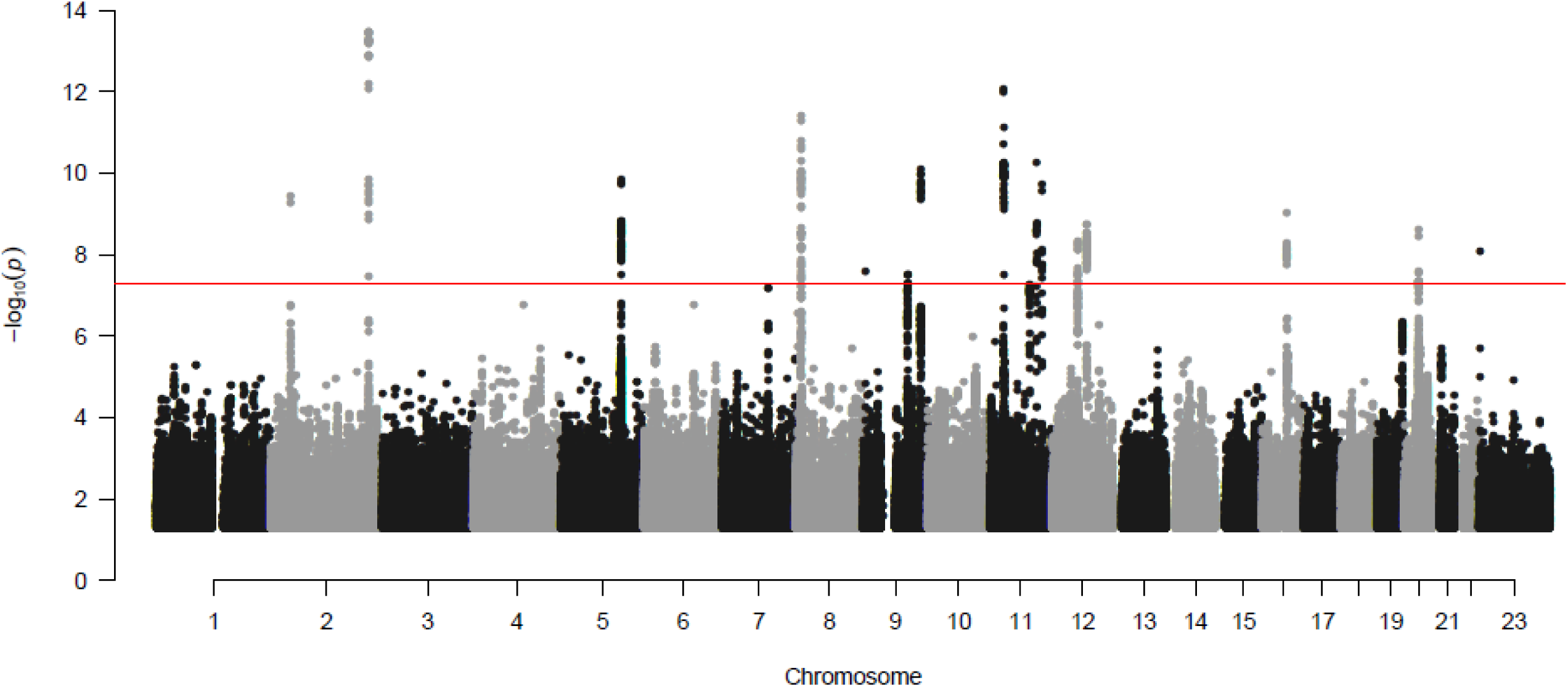
Manhattan plot indicating genome-wide significant variants. The X axis indicates the chromosome depicted by alternating black and grey. The Y axis indicates the inverse log10 of the p-value (-log10(p)). The line designates the minimum p-value for genome-wide significance.

**Figure 2.**
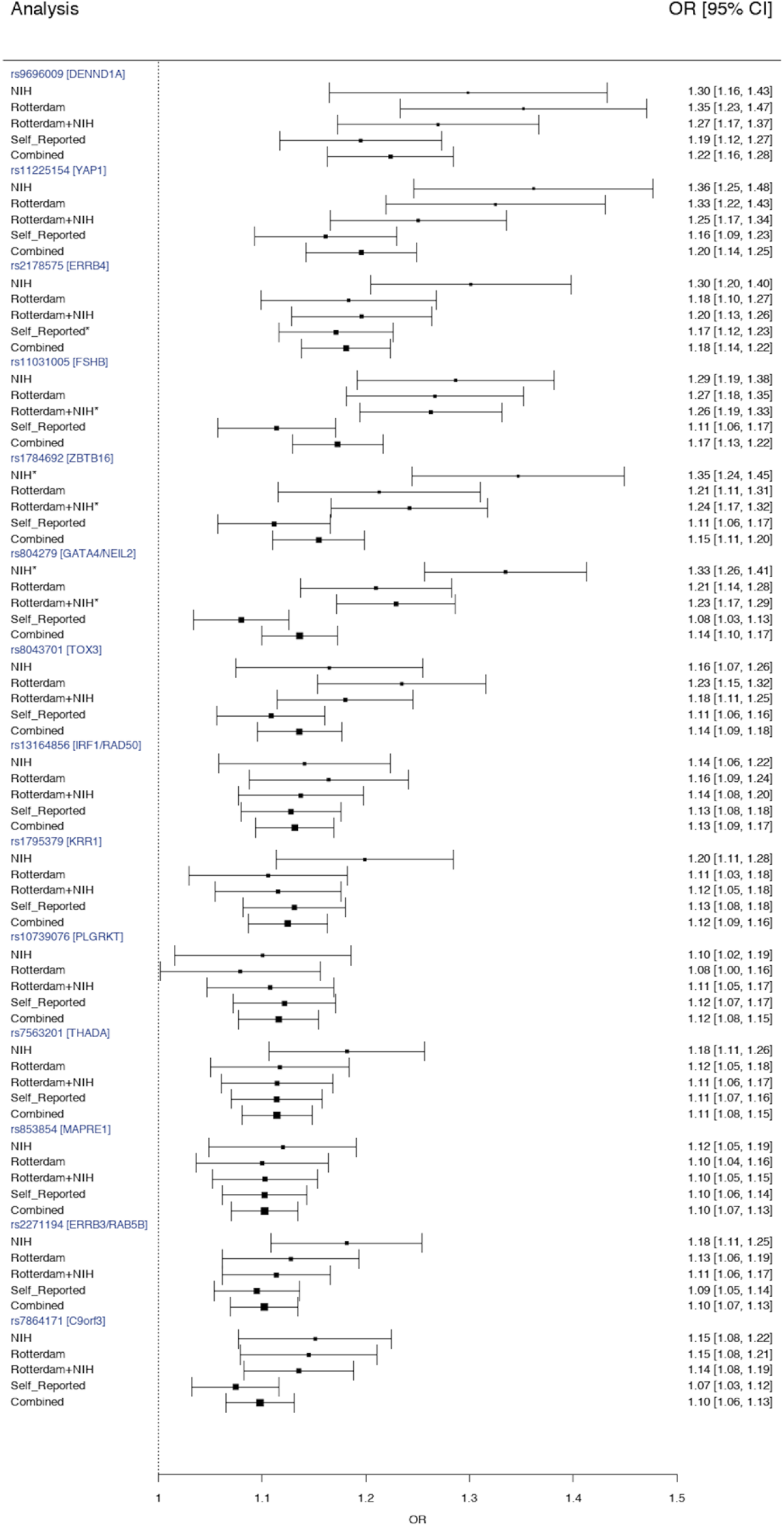
Odds ratio of polycystic ovary syndrome (PCOS) as a function of diagnostic criteria applied. The Y-axis specifies the diagnostic criteria and the X-axis indicates the odds ratio (OR) and 95% confidence intervals (CI) for PCOS (black circle and horizontal error bars). Data derived as follows: NIH=groups recruiting only NIH diagnostic criteria; Rotterdam=Rotterdam groups recruiting Rotterdam diagnostic criteria including the subset fulfilling NIH diagnostic criteria; Rotterdam+NIH=all groups except self-reported; self-reported=23andME; and combined= all groups. Specific OR’s [95% CI, 5% CI] are indicated on the right. rs804279 in the GATA4/NEIL2 locus demonstrates significant heterogeneity (Het P =2.9×10^-4^). The * indicates statistically significant association for PCOS and the variant in that specific stratum.

**Table 2.**
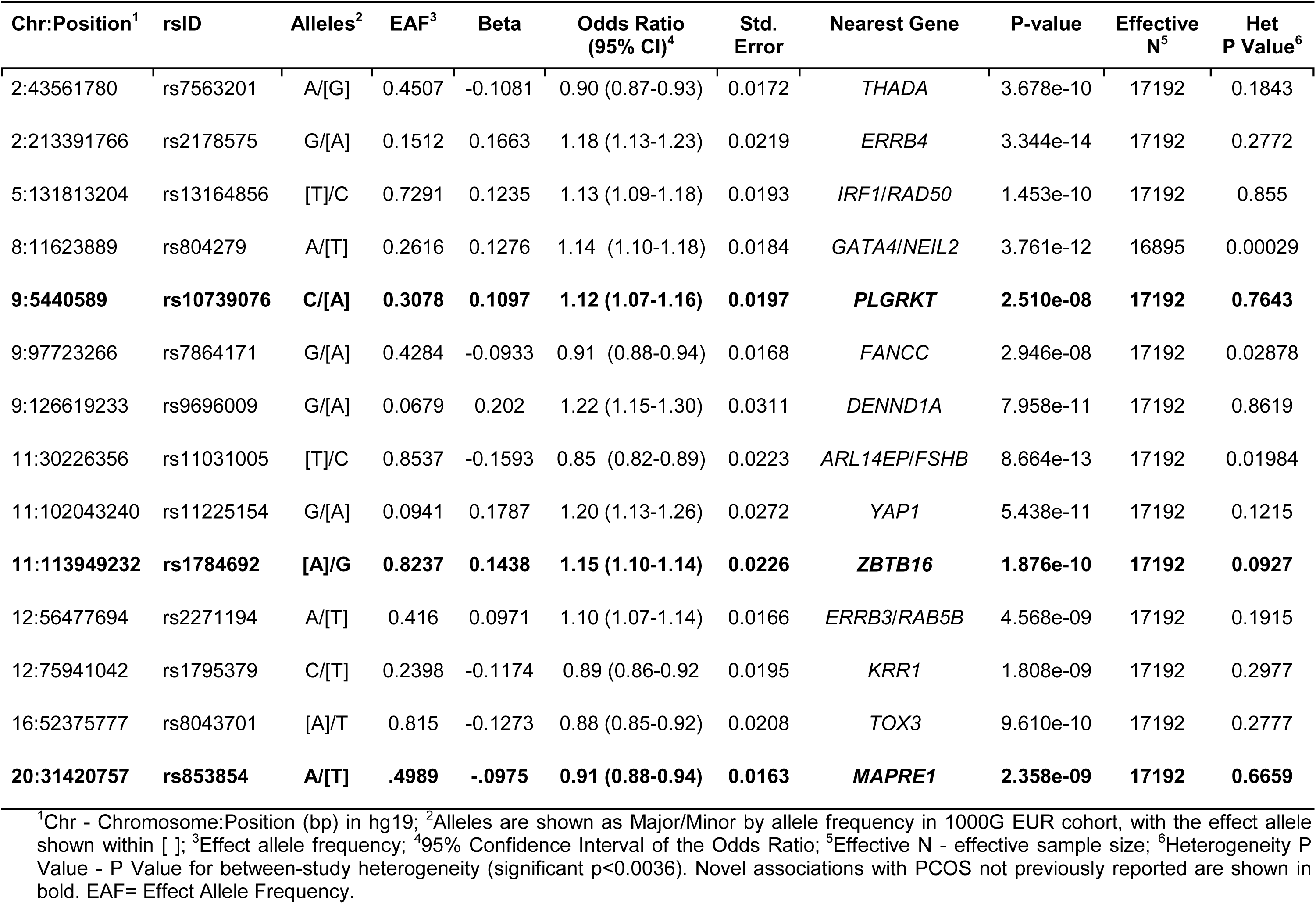
The 14 genome-wide significant variants associated with PCOS in the meta-analysis.

We assessed the association of the PCOS susceptibility variants identified in the GWAS meta-analysis with the PCOS related traits: HA, OD, PCOM, testosterone, FSH and LH levels, and ovarian volume in PCOS cases (**Table 3**, **Supplementary Figure 2** and **Supplementary Table 4**). We found four variants associated with HA, eight variants associated with PCOM and nine variants associated with OD. Of the eight loci associated with PCOM, seven were also associated with OD. Three of the four loci associated with HA were also associated with OD and PCOM. Two additional loci were associated with OD alone, one of which was the locus near *FSHB* (**Supplementary Table 4**). This locus was also associated with LH and FSH levels. There was a single PCOS locus near *IRF1/RAD50* associated with testosterone levels (**Supplementary Table 4**). We repeated this analysis with susceptibility variants reported previously in Han Chinese PCOS cohorts^14, 15^. In this analysis, there was one association with HA (near *DENND1A*), three with PCOM and three with OD (**Supplementary Table 3**). A limitation of these analyses is the variable sample size across the phenotypes analysed. Additionally, the known referral bias for the more severely affected NIH phenotype (patients having both OD and HA) may result in more PCOS diagnoses than the other criteria^35^, and may have contributed to the number of associations between the identified PCOS risk loci and these phenotypes.

**Table 3.**
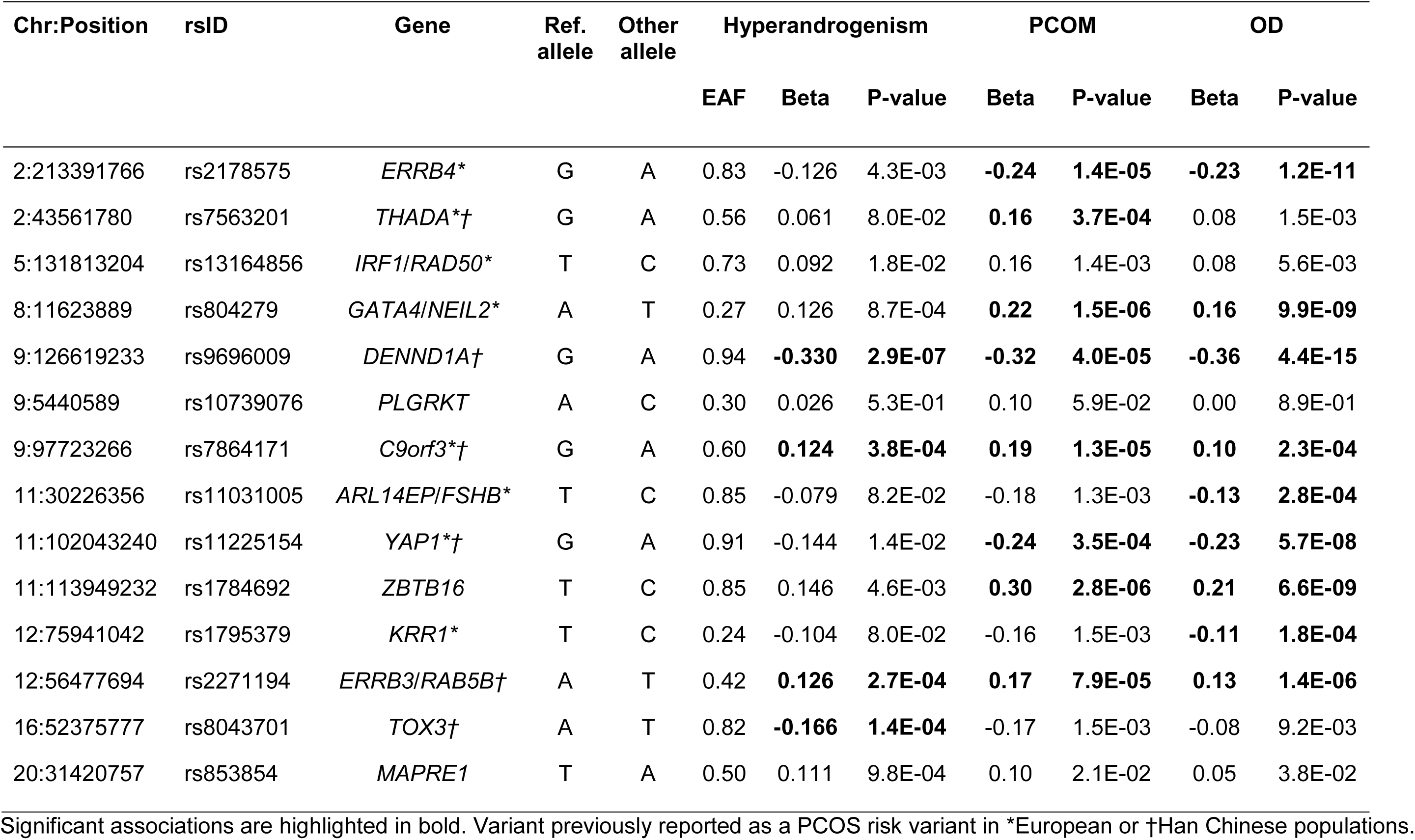
Association of PCOS GWAS Meta-analysis Susceptibility Variants and PCOS related traits.

In the analyses looking at the weighted genetic risk score in the Rotterdam cohort, we observed an increase in the risk for PCOS (**Supplementary Figure 3**). Compared to individuals in the third quintile (reference group), individuals in the top 5th quintile of risk score have an OR of 2.78 (95% CI) for PCOS based on NIH criteria and an OR of 3.39 (95% CI) for Rotterdam criteria based PCOS. Of the associations, only the effect estimate for the Rotterdam criteria was significant, possibly due to the smaller size available with cases diagnosed according to the NIH criteria. When looking at the area under the ROC curves at SNPs with different P-value thresholds, we found a maximum AUC of 0.54 using SNPs with a P-value < 5×10^-6^ for both diagnostic criteria. While this is significantly better than chance, it is unlikely that a risk score generated from the variants discovered to date would represent a clinically relevant tool.

**Table 4.** LD Score regression results using the LDSC method.

| Phenotype | Genetic Correlation | SE | Z | P-value |
| --- | --- | --- | --- | --- |
| Body mass index | 0.34 | 0.039 | 8.60 | $8.21 \times 10^{-18}$ |
| Childhood obesity | 0.34 | 0.066 | 5.17 | $2.40 \times 10^{-7}$ |
| Fasting insulin levels | 0.44 | 0.087 | 5.01 | $5.33 \times 10^{-7}$ |
| Type 2 diabetes | 0.31 | 0.068 | 4.47 | $7.84 \times 10^{-6}$ |
| High-density lipoprotein levels | -0.23 | 0.059 | -3.96 | $7.40 \times 10^{-5}$ |
| Menarche | -0.16 | 0.042 | -3.76 | $1.71 \times 10^{-4}$ |
| Triglyceride levels | 0.19 | 0.052 | 3.61 | $3.05 \times 10^{-4}$ |
| Coronary artery disease | 0.23 | 0.069 | 3.32 | $8.86 \times 10^{-4}$ |
| Depression | 0.205 | 0.0582 | 3.5203 | 0.0004 |
| Menopause | -0.014 | 0.0183 | -0.762 | 0.4461 |
| Male pattern balding | 0.0149 | 0.0168 | 0.8861 | 0.3756 |

LD score regression analysis revealed genetic correlations with childhood obesity, fasting insulin, T2D, HDL, menarche timing, triglyceride levels, cardiovascular diseases and depression (**Table 4**) suggesting that there is shared genetic architecture and biology between these phenotypes and PCOS. There were no genetic correlations with menopause timing or male pattern balding. Mendelian randomization suggested that there was a causal role for BMI, fasting insulin and depression pathways (**Table 5**). Interestingly, while there was no genetic correlation detected for male pattern balding or menopause timing with PCOS, the Mendelian randomization analysis was significant. The importance of BMI pathways on reproductive phenotypes was further demonstrated by the attenuation of significance of Mendelian randomization analysis for age-at-menarche when BMI-associated variants were excluded from the analysis.

**Table 5.** Mendelian randomization using an inverse weighted variant method.

| Potential Risk factor | IVW method |  |  |
| --- | --- | --- | --- |
|  | Beta | SE | P-value |
| Body mass index | 0.72 | 0.072 | $1.56 \times 10^{-23}$ |
| Fasting insulin levels* | 0.03 | 0.007 | $1.73 \times 10^{-5}$ |
| Male pattern balding | 0.05 | 0.017 | 0.0034 |
| Menopause | 0.1 | 0.022 | $1.31 \times 10^{-5}$ |
| Depression | 0.77 | 0.213 | 0.00029 |

\*Loci used were initially reported in an analysis of fasting insulin adjusted for BMI. IVW = inverse weighted variant.

We found 14 independent loci significantly associated with the risk for PCOS, including three novel loci. The 11 previously reported loci implicated neuroendocrine and metabolic pathways that may contribute to PCOS (**Supplementary Note 1.1**). Two of the novel loci contain potential endocrine related candidate genes. The locus harbouring rs10739076 contains several interesting candidate genes; *PLGRKT,* a plasminogen receptor and several genes in the insulin superfamily; *INSL6, INSL4* and *RLN1, RLN2* which are endocrine hormones secreted by the ovary and testis and are suspected to impact follicle growth and ovulation^22^. *ZBTB16* (also known as *PLZF*) has been marked as an androgen-responsive gene with anti-proliferative activity in prostate cancer cells^23^. *PLZF* activates *GATA4* gene transcription and mediates cardiac hypertrophic signalling from the angiotensin II receptor 2^24^. Furthermore, *PLZF* is upregulated during adipocyte differentiation *in vitro*^25^ and is involved in control of early stages of spermatogenesis^26^ and endometrial stromal cell decidualization^27^. The third novel locus harbours a metabolic candidate gene; *MAPRE1* (interacts with the low-density lipoprotein receptor related protein 1 (LRP1), which controls adipogenesis^28^ and may additionally mediate ovarian angiogenesis and follicle development^29^ (**Supplementary Note 1.2**). Thus, all the new loci contain genes plausibly linked to both the metabolic and reproductive features of PCOS.

We found that there was no significant difference in the association with case status for the majority of the PCOS-susceptibility loci by diagnostic criteria. However, due to lack of power, it was not possible to compare each of the four PCOS phenotypes (Supplementary Figure 1). Cohorts containing both NIH and non-NIH Rotterdam phenotypes were compared to cohorts containing only the NIH phenotype. Although the inclusion of NIH phenotypes in the Rotterdam criteria cohorts could have masked differences in genetic architecture between the NIH and non-NIH Rotterdam phenotypes, the non-NIH Rotterdam subjects made up 53% of the cohorts. It is of considerable interest that the cohort of research participants from the personal genetics company 23andMe, Inc., identified by self-report, had similar risks to the other cohorts where the diagnosis was clinically confirmed. Our findings suggest that the genetic architecture of these PCOS definitions does not differ for common susceptibility variants. If these results are confirmed when the Rotterdam diagnostic subsets are compared (**Supplementary Figure 1**), data should be combined in future efforts to map the genetic architecture and molecular pathways underlying PCOS. Only one locus, *GATA4/NEIL2* (rs804279), was significantly different across diagnostic criteria: most strongly associated in NIH compared to the Rotterdam phenotype and self-reported cases. Deletion of *GATA4* results in abnormal responses to exogenous gonadotropins and impaired fertility in mice^30^. The locus also encompasses the promoter region of *FDFT1,* the first enzyme in the cholesterol biosynthesis pathway^31^, which is the substrate for testosterone synthesis, and is associated with non-alcoholic fatty liver disease^32^. The major difference between the NIH phenotype and the additional Rotterdam phenotypes is metabolic risk; the NIH phenotype is associated with more severe insulin resistance^33^. rs804279 does not show association with any of the metabolic phenotypes in the T2D diabetes knowledge portal {Type 2 Diabetes Knowledge Portal. type2diabetesgenetics.org. 2015 Feb 1; http://www.type2diabetesgenetics.org/variantInfo/variantInfo/rs804279} so it may represent a PCOS-specific susceptibility locus.

The significant association of PCOS GWAS meta-analysis susceptibility variants with the cardinal PCOS related traits OD, HA and PCOM further strengthened the hypothesis that specific variants may confer risk for PCOS through distinct mechanisms. Three variants at the *C9orf3, DENND1A,* and *RAB5B* were associated with all PCOS related traits. The findings were consistent with the Han Chinese *DENND1A* variant association with HA, as suggested previously^34^. Thus, these loci, along with *GATA4/NEIL2 (as discussed above*) may help identify pathways that link specific PCOS related traits with greater metabolic risk. In contrast, the variants at the *ERBB4, YAP1,* and *ZBTB16* loci were strongly associated with OD and PCOM, and therefore, might be more important for links to menstrual cycle regularity and fertility. In addition, the *FSHB* variant was associated with the levels of FSH and LH^16, 17^, suggesting that it may act by affecting gonadotropin levels. This variant maps 2kb upstream from open chromatin (identified by DNase-Seq) and an enhancer (identified by peaks for both H3K27Ac and H3K4me1) in a lymphoblastoid cell line from ENCODE, indicating a potential role for a regulatory element ∼25kb upstream from the *FSHB* promoter. Furthermore, the association between the *IRF1/RAD50* variant and testosterone levels may indicate a regulatory role in testosterone production.

Of note, results of the follow-up analysis show a high level of shared biology between PCOS and a range of metabolic outcomes consistent with the previous findings^17^. In particular, there is genetic evidence for increased BMI as a risk factor for PCOS. There is also genetic evidence that fasting insulin might be an independent risk factor, but there might also be pleiotropic effects across the variants given the results from different Mendelian randomization analyses. This study also confirmed a causal association with the pathways that underlie menopause^17^, suggesting that PCOS has shared aetiology with both classic metabolic and reproductive phenotypes. Furthermore, there was an apparent effect of depression-associated variants on the likelihood of PCOS, suggesting a role for psychological factors on hormonally related diseases. However, the links between PCOS and depression might be complicated by pathways that are also related to BMI. In addition, male-pattern balding-associated variants showed strong effects on PCOS, suggesting that this might be a male manifestation of PCOS pathways, as has been previously suggested^18, 20, 21, 36^. This observation may reflect the biology of hair follicle sensitivity to androgens, seen in androgenetic alopecia, a well-recognised feature of HA and PCOS ^37, 38^. The Mendelian randomization results for male-pattern balding and menopause are significant despite non-significant genetic correlation results, suggesting that the shared aetiology may be specific to only a few key pathways.

In conclusion, the genetic underpinnings of PCOS implicate neuroendocrine, metabolic and reproductive pathways in the pathogenesis of disease. Although specific phenotype stratified analyses are needed, genetic findings were consistent across the diagnostic criteria for all but one susceptibility locus, suggesting a common genetic architecture underlying the different phenotypes. There was genetic evidence for shared biologic pathways between PCOS and a number of metabolic disorders, menopause, depression and male-pattern balding, a putative male phenotype. Our findings demonstrate the extensive power of genetic and genomic approaches to elucidate the pathophysiology of PCOS.

## ACKNOWLEDGEMENTS

We thank the research participants and employees of 23andMe for contributing to this study. EGCUT Computations were performed in High Performance Computing Center, University of Tartu.

## ACKOWLEDGEMENT OF FUNDING SOURCES

This work has been supported by; MRC grant MC_U106179472 (F.D.), Samuel Oschin Comprehensive Cancer Institute Developmental Funds, Center for Bioinformatics and Functional Genomics and Department of Biomedical Sciences Developmental Funds (M.R.J.), NCI P30CA177558 (C.H.), NCI UM1CA186107 (P.K.),

European Regional Development Fund (Project No. 2014-2020.4.01.15-0012) and the European Union’s Horizon 2020 research and innovation program under grant agreements No 692065 (T.L., R.M. A.S.) and 692145 (R.M.), NICHD R01HD065029 (R.S.), Estonian Ministry of Education and Research (grant IUT34-16) (T.L.), NICHD R01HD057450, P50HD044405 (M.U., M.G.H, A.D.), NICHD R01HD057223, R01HD085227 (M.G.H. A.D.), deCode Genetics (G.T. U.T., K.S., U.S.), Raine Medical Research Foundation Priming Grant (B.H.M.), NIHR BRC, Wellcome Trust, MRC (T.S.), NIDDK U01DK094431, U01DK048381 (D.E.), NICHD U10HD38992 (R.L.), Estonian Ministry of Education and Research (grant IUT34-16), Enterprise Estonia (grant EU48695); the EU-FP7 Marie Curie Industry-Academia Partnerships and Pathways (IAPP, grant SARM, EU324509) (A.S.), Wellcome (090532, 098381, 203141); European Commission (ENGAGE: HEALTH-F4-2007-201413) (M.McC.), MRC G0802782, MR/M012638/1 (S.F.), NICHD R01HD065029, ADA 1-10-CT-57, Harvard Clinical and Translational Science Center, from the National Center for Research Resources 1UL1 RR025758 (C.W.).

## ADDITIONAL AUTHOR INFORMATION 23andMe RESEARCH TEAM

Michelle Agee, Babak Alipanahi, Adam Auton, Robert K. Bell, Katarzyna Bryc, Sarah L. Elson, Pierre Fontanillas, Nicholas A. Furlotte, David A. Hinds, Karen E. Huber, Aaron Kleinman, Nadia K. Litterman, Matthew H. McIntyre, Joanna L. Mountain, Elizabeth S. Noblin, Carrie A.M. Northover, Steven J. Pitts, J. Fah Sathirapongsasuti, Olga V. Sazonova, Janie F. Shelton, Suyash Shringarpure, Chao Tian, Joyce Y. Tung, Vladimir Vacic, Catherine H. Wilson.

## COI statement

Members of the 23andMe Research team are employees of and hold stock or stock options in 23andMe, Inc. G.T., U.T., K.S., U.S., are employees of deCODE genetics/Amgen Inc. MMcC serves on advisory panels for Pfizer and NovoNordisk; has received honoraria from Pfizer, NovoNordisk and EliLilly; and received research funding from Pfizer, NovoNordisk, EliLilly, AstraZeneca, Sanofi Aventis, Boehringer Ingelheim, Merck, Roche, Janssen, Takeda, Servier. J.L. has received consultancy fees from Danone, Metagenics inc., Titus Healthcare, Roche and Euroscreen. C.W. is a consultant for Novartis and has received UptoDate royalties.

## ONLINE METHODS

### Methods

#### Subjects

The meta-analysis included 10,074 cases and 103,164 controls from seven cohorts of European descent. For the analysis of PCOS related traits three additional cohorts, the Northern Finnish Birth Cohort (NFB66)^39^, Twins UK^40^ and the Nurses’ Health Study (NHS)^41^ were included. Cases were diagnosed with PCOS based on NIH or Rotterdam Criteria or by self-report. The NIH criteria require the presence of both OD and clinical and/or biochemical HA for a diagnosis of PCOS^6^. The Rotterdam criteria require two out of three features 1) OD defined by oligo- or amenorrhea, 2) clinical and/or biochemical hyperandrogenism (HA) and/or 3) PCOM for a diagnosis of PCOS^7^. Self-reported cases from research participants in the 23andMe, Inc. (Mountain View, CA, USA) cohort either responded “yes” to the question “Have you ever been diagnosed with polycystic ovary syndrome?” or indicated a diagnosis of PCOS when asked about fertility (“Have you ever been diagnosed with PCOS?” or “What was your diagnosis? Please check all that apply.” Answer=PCOS), hair loss in men or women (“Have you been diagnosed with any of the following? Please check all that apply.” Answer=PCOS) or research question (“Have you ever been diagnosed with PCOS?”)^17^.

HA was defined as hirsutism and quantified by the Ferriman-Gallwey (FG) score. The FG score assesses terminal hair growth in a male pattern in females, and a score above the upper limit of normal controls (>8) is considered hirsutism^42^. Hyperandrogenemia was defined as testosterone, androstenedione or DHEAS greater than the 95% confidence limits in control subjects in the individual population. OD was defined as cycle interval <21 or >35 days^43^. PCOM was defined as 12 or more follicles of 2-9 mm in at least one ovary or an ovarian volume >10 mL^7^. The quantitative PCOS traits included levels of total testosterone (T), follicle-stimulating hormone (FSH), and luteinizing hormone (LH) and ovarian volume (Supplementary Table 1). An overview of the cohorts, diagnostic criteria and number of subjects included are summarized in Table 1 and Supplementary Table 1. All studies were approved by the Institutional Review Board at the recruiting site and all subjects signed written, informed consent.

#### Data Collection and Quality Control

Each study provided summary results of genetic per-variant-estimates produced in either case-control or trait association analyses. The collected files underwent central quality control (QC) using the EasyQC pipeline^44^. Variants were excluded based on minor allele frequency (MAF) < 1%, imputation quality (R^2^) < 0.3 or info < 0.4 for MACH and IMPUTE2 respectively^45, 46^.

#### Meta-analysis of PCOS status and PCOS related traits

The per-variant-estimates collected from the contributing studies were meta-analyzed using a fixed-effect, inverse-weighted-variance meta-analysis that employed either GWAMA^47^ or METAL^48^. In addition to the overall meta-analysis, we performed meta-analyses for studies with available data for the separate PCOS diagnostic criteria: NIH, Rotterdam^7^ and self-report^17^, as well as for the PCOS related traits of HA, OD and PCOM. The meta-analysis of PCOS status was performed using two models; (1) age-adjusted, (2) age and BMI-adjusted, given the high prevalence of obesity in affected women that resulted in cases being significantly heavier than controls in most cohorts (Table 1).

We removed any variants that were not present in more than 50% of the effective sample size prior to combining with 23andMe as this was the largest cohort in the meta-analysis, providing approximately 51% of the PCOS cases and 80% of controls. We also removed any variants only present in one study. The meta-analysis of PCOS related traits was performed adjusting for age and BMI. Identified variants were annotated for insight into their biological function using ANNOVAR^49^ to assign refGene gene information, SIFT score^50^, PolyPhen2 scores^51^, CADD scores^52^, GERP scores^53^ and SiPhy log odds^54^.

#### Comparison of PCOS Diagnostic Criteria

Since the Rotterdam diagnostic criteria include the NIH phenotype of OD and HA, the Rotterdam criteria cohorts included a subset of NIH phenotype cases, as indicated in Table 1. The Rotterdam criteria cohorts containing the NIH phenotype were compared to the NIH criteria cohorts for the analysis of the genetic architecture of the diagnostic criteria. We lacked adequate statistical power to compare the four PCOS phenotypes (Supplementary Figure 1) or to compare the NIH to the non-NIH Rotterdam phenotypes.

#### Identifying Associations Between PCOS Loci and PCOS related traits

In order to understand biology relevant to identified PCOS susceptibility, we assessed the association between index SNPs at each genome-wide-significant locus and the PCOS related traits HA, OD, PCOM as well as the quantitative traits testosterone, LH and FSH levels and ovarian volume. The threshold for significance in this analysis was p<4.5×10^-4^ (Bonferroni correction [0.05/(14 independent loci x 8 traits)].

#### Identifying Shared Risk Loci Between European Ancestry and Han Chinese PCOS

In order to identify shared risk loci between the previously reported GWAS in Han Chinese PCOS cases and our European ancestry cohort, 13 independent signals (represented by 15 SNPs) at 11 genome-wide significant loci reported by Chen *et al.*^14^ and Shi *et al.*^15^ were investigated for association in our meta-analyses of PCOS and PCOS related traits. The adjusted P-value for this analysis was <0.00048 (Bonferroni correction [0.05/(13 independent signals x 8 traits)]).

#### Biologic Function of Genes in Associated Loci

Information on the biological function of the nearest gene (or genes, if variants were equidistant from more than one coding transcript and annotated as such by ANNOVAR^49^ to the index SNP of each identified risk locus) was collected by performing a search of the Entrez Gene Database^55^, and collecting the co-ordinates of the gene (genome build 37; hg19) as well as the cytogenetic location and the summary of the gene function. In addition to the EntrezGene Database queries, the gene symbol was used as a search term in the PubMed database^56^, either alone or combined with the additional search term “PCOS” to identify relevant published literature in order to obtain information on putative biological function and involvement in the pathogenesis of PCOS (summarized in Supplementary Text Section 1.1).

#### Weighted genetic risk score and prediction

One potential use of genetic risk scores is prediction of disease. The ability of genetic risk scores calculated from loci discovered in analysis of the different diagnostic criteria to discriminate cases from alternative criteria was measured. We constructed a weighted genetic risk score based on a meta-analysis excluding the Rotterdam Study subjects. The weighted genetic risk score was divided into quintiles and tested for association with PCOS in the Rotterdam cohort. The middle quintile was used as the reference and the odds for having PCOS based on both Rotterdam or NIH criteria was then calculated.

Additionally, the 23andMe results were used to select independent SNPs with cut-offs of *p*<5×10^-4^ to *p*<5×10^-8^. The Rotterdam cohort was then used to calculate risk scores and the area-under-the curve (AUC) for both NIH and Rotterdam diagnostic criteria. Analyses were performed using PLINK v1.9 and SPSS v21 (IBM Corp, Armonk, NY)^57^

#### Linkage Disequilibrium (LD) Score Regression

To assess the level of shared etiology between PCOS and related traits, we performed genetic correlation analysis using LD-score regression ^58^. Publicly available genome-wide summary statistics for body mass index (BMI) ^59^, childhood obesity ^60^, fasting insulin levels (adjusted for BMI) ^61^, type 2 diabetes ^62^, high-density lipoprotein (HDL) levels ^63^, menarche timing ^64^, triglyceride levels ^63^, coronary artery disease ^65^, depression ^63^, menopause^17^ and male pattern balding ^66^ were used to estimate the genome-wide genetic correlation with PCOS. The adjusted P-value for this analysis was p<0.0045 after a Bonferroni correction (0.05/11 traits).

#### Mendelian Randomization

Phenotypes of interest, both where there was evidence of shared genetic architecture and where there was previous evidence for genetic links, were assessed using Mendelian randomization methods. Mendelian randomization differs from LD score regression in that one phenotype is analysed as a potential causal factor for another. Mendelian randomization was performed using both inverse weighted variance and Egger’s regression methods ^67^, with inverse weighted methods being more powerful, but Egger’s methods being resistant to directional pleiotropy (where there are a set of SNPs that appear to have an alternative pathway of effect). In addition to the phenotypes implicated by the LD-score regression measures, male pattern balding has a strong biological rationale and was therefore included. The genetic score for childhood obesity substantially overlaps with the score for adult BMI (such that the INSIDE violation - where the effect of SNPs on a confounding factor scales with that on the trait of interest - of Mendelian randomization would likely occur^68^, so only a score for BMI was used, with the proviso that this represents BMI across the whole of the life course after very early infancy. The SNPs for Depression were drawn from the results of a more recent analysis, for which there was not, at time of analysis, publicly available genome-wide data.

#### Credible Sets

We defined a locus as mapping within 500kb of the lead SNP. For each locus, we first calculated the posterior probability, π_Cj_, that the jth variant is driving the association, given byc

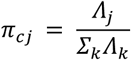

where the summation is over all retained variants in the locus. In this expression, Λ_j_ is the approximate Bayes’ factor ^69^ for the jth variant, given by

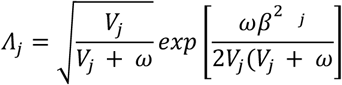

where β_j_ and V_j_ denote the estimated allelic effect (log-OR) and corresponding variance from the meta-analysis. The parameter ω denotes the prior variance in allelic effects, taken here to be 0.04 ^69^. The 99% credible set ^70^ for each signal was then constructed by: (i) ranking all variants according to their Bayes’ factor, Λ_j_; and (ii) including ranked variants until their cumulative posterior probability of driving the association attained or exceeded 0.99.

## SUPPLEMENTARY MATERIALS

### SUPPLEMENTARY NOTES

#### 1.1. Supplementary Results

In addition to these 14 significant loci, there was suggestive evidence of a 15th signal, rs151212108, near *ARSD* on the X chromosome. This SNP shows a relatively large effect size (OR:1.72, CI:1.43-2.07, P=8.35×10^-9^). However, the SNP had low sample number (overall minor allele frequency=0.0765) with poor imputation quality and it was present in only three studies; Oxford, deCODE, and Chicago. Further, the signal showed nominally significant heterogeneity (P =0.028) in the direction of effect estimates between Oxford, where the effect allele had a protective effect, and deCODE and Chicago, where the effect allele increased risk of PCOS. Thus, this signal was less robust than our other signals and will require further confirmation. Accordingly, we have not included this locus in downstream analyses. A detailed review of genes within reported loci is included in the Supplementary Notes (Section 1.2).

#### 1.2 Literature Lookup of genes at PCOS risk loci

Summary of published literature on gene function of PCOS susceptibility loci.

1. **THADA (Thyroid Adenoma Associated); Located at 2p21 (Chr 2: 43561780-43561780).** Encodes a transcript of largely unknown function. THADA encodes thyroid adenoma-associated protein, which is expressed in pancreas, adrenal medulla, thyroid, adrenal cortex, testis, thymus, small intestine, and stomach^71^. This gene has been identified in GWAS for gestational weight gain, inflammatory bowel disease and PCOS and more specifically with the phenotype trait PCOM ^72,73,73,14, 71^. THADA has been associated with endocrine and metabolic disturbances commonly found in PCOS, such as increased LH, testosterone and LDL levels and T2D^75^. Proposed to modify PCOS risk through metabolic mechanisms ^76^.
2. **ERBB4 (erb-b2 receptor tyrosine kinase 4; also known as HER4); Located at 2q33.3-q34.** Fourth member of the EGFR (epidermal growth factor *receptor*) family and the Tyr protein kinase family (USCS, GeneNetwork, RefSeq). Participates in the YAP/Hippo pathway, which regulates cell proliferation, differentiation and apoptosis and has been associated with the size of the primordial follicle pool in mice, female reproductive capacity in Drosophila, and it is hypothesized that disruption of Hippo signaling can promote follicle growth^77–81^. Tyrosine-protein kinase plays an essential role as cell surface receptor for neuregulins and EGF family members and regulates development of the heart, the central nervous system and the mammary gland ^82^. HER4 is characterized by anti-proliferative and pro-apoptotic activity, is co-expressed in 90% of ER positive breast tumors. Proposed to modify PCOS risk through metabolic mechanisms ^76^. Suggested to have a pathogenic role in cystogenesis in polycystic kidney disease ^83^.
3. **IRF1 (interferon regulatory factor 1): Located at 5q31.1 (Chr5: 131813204-131813204);** Belongs to the interferon regulatory transcription factor (IRF) family. Activates the transcription of interferons alpha and beta, and genes induced by interferons alpha, beta and gamma (USCS, GeneNetwork, NCBI gene database). IRF1 displays a functional diversity in the regulation of cellular responses including host response to viral and bacterial infections, inflammation, and cell proliferation and differentiation, regulation of the cell cycle and induction of growth arrest and programmed cell death following DNA damage (UniProtKB). Acts as a tumor suppressor and plays a role not only in antagonism of tumor cell growth but also in stimulating an immune response against tumor cells^84^. It has been shown in fathead minnow that IRF1 may function as early molecular switches to control phenotypic changes in ovary tissue architecture and function in response to androgen or antiandrogen exposure^85^.
4. **RAD50 (RAD50 homolog): Located at 5q31. (Chr5: 131892616-131980313)**; Rad50, a protein involved in DNA double-strand break repair^82^. This protein forms a complex with MRE11 and NBS1. The protein complex binds to DNA and displays numerous enzymatic activities that are required for non-homologous joining of DNA ends, and is important for DNA double-strand break repair^86^, telomere maintenance, and meiotic recombination^87^. Knockout studies of the mouse homolog suggest this gene is essential for cell growth and viability^88^.
5. **GATA4 (GATA binding protein 4): Located at 8p23.1-p22 (Chr 8: 11623889-11623889).** GATA4 encodes a zinc-finger transcription factor that recognizes the GATA motif in the promoters of various genes (RefSeq). GATA4 is implicated in regulating granulosa cell differentiation, proliferation and function and is expressed in follicles, embryoid bodies and chorion of women with PCOS^89^. Knockdown of GATA4 and GATA6 impairs folliculogenesis and induces infertility^30, 90^. The loss of GATA4 within the ovary results in impaired granulosa cell proliferation and theca cell recruitment^91^. Knockdown of both genes affects expression of FSH receptor, LH receptor, inhibin α and β^30, 89^. In rats with reproductive and metabolic abnormalities similar to PCOS, GATA4 has been associated with the biosynthesis and metabolism of steroids^92^. It is also proposed to modify PCOS risk through metabolic and inflammatory mechanisms ^76^.
6. **PLGRKT (Plasminogen receptor, C-terminal lysine transmembrane protein): Located at 9p24.1. (Chr 9; 5440339-5440839).** PLGRKT encodes a plasminogen receptor involved in regulating macrophage migration and regulates catecholamine release^93^. The region also includes genes for several members of the insulin superfamily (INSL6, INSL4, RLN1, RLN2), which have roles in spermatogenesis, follicle growth and ovulation^22, 94^.
7. **FANCC (Fanconi anemia, complementation group C): Located at: 9q22.3 (Chr 9: 97723266-97723266).** Member of the Fanconi anemia complementation group which, amongst others, includes FANCD1 (BRCA2). Members of the Fanconi anemia complementation group are related by their assembly into a common nuclear protein complex (RefSeq). This gene encodes the protein for complementation group C. Fanconi anemia is a recessive repair deficiency disorder, characterized by cytogenetic instability, hypersensitivity to DNA crosslinking agents, chromosomal breakage and defective DNA repair (UCSC, GeneNetwork). FANCC is a DNA repair protein that may operate in a post replication repair or a cell cycle checkpoint function^82^. May be implicated in interstrand DNA cross-link repair and in the maintenance of normal chromosome stability. Was recently shown to have a mitophagy function as well, and is required for clearance of damaged mitochondria ^95^.
8. **C9orf3. Located at 9q22.32.** Has been previously associated with PCOS^15^. However, the region also includes genes for two hormones that regulate gluconeogenesis (FBP1, FBP2), and for PTCH1, which is a receptor for hedgehog proteins. In mice, the hedgehog signaling has been shown to be important for ovarian follicle development and is also implicated in the proliferation and steroidogenesis of theca cells^96^. This is supported by the association between rs4385527 in C9orf3 and anovulation, HA and polycystic ovarian morphology (PCOM)^73^.
9. **DENND1A (DENN/MADD domain-containing protein 1A): Located at 9q33.3 (Chr 9: 126619233-126619233).** Member of the connecdenn family and functions as a guanine nucleotide exchange factor involved for the early endosomal small GTPase RAB35 (UCSC, RefSeq). Regulates clathrin-mediated endocytosis (a major mechanism for internalization of proteins and lipids) through RAB35 activation (USCS, RefSeq). DENND1A variant 2 (DENND1A.V2) protein and mRNA levels are increased in PCOS theca cells and play a key role in the hyperandrogenemia associated with PCOS^97^. The *DENND1A* locus has also been associated with PCOM and elevated serum insulin levels in PCOS women^73, 75^. Some SNP’s in DENND1A have even been associated with endometrioid carcinoma. It has been suggested that *DENND1A, LHCGR, INSR*, and *RAB5B* form a hierarchical signalling network that can influence androgen synthesis^98^.
10. **ARL14EP (ADP-ribosylation factor-like 14 effector protein): Located at 11p14.1.** Encodes an effector protein, which interacts with ADP-ribosylation factor-like 14 [ARL14], beta-actin and actin-based motor protein myosin 1E. ARL14 controls the export of major histocompatibility class II molecules by connecting to the actin network via this effector protein (RefSeq).
11. **FSHB (Follicle stimulating hormone, beta polypeptide): Located at 11p14.1 (Chr 11; 30226356-30226356).** FSHB is a member of the pituitary glycoprotein hormone family and encodes the β-subunit of the follicle-stimulating hormone (FSH) (RefSeq). FSH regulates folliculogenesis. FSHB polymorphisms influence early follicular phase FSH concentrations and IVF treatment outcome^99^. SNPs in the FSHB region are known to be associated with circulating FSH, LH and AMH levels but also with PCOS ^99–105^. Overexpression of FSHB could cause polycystic ovary syndrome in women, whereas inactivating mutations of the FSHB gene, encoding for the hormone’s unique β-subunit, cause infertility by primary amenorrhea ^106, 107^.
12. **YAP1 (Yes-associated protein 1): Located at 11q13 (Chr 11; 102043240-102043240).** YAP1 is an effector protein in the Hippo pathway involved in development, growth, repair, and homeostasis (RefSeq). This pathway also plays a pivotal role in organ size control and tumor suppression by restricting proliferation and promoting apoptosis^82^. Has been associated with the size of the primordial follicle pool in mice, female reproductive capacity in *Drosophila*, and it is hypothesized that disruption of Hippo signaling can promote follicle growth^79^. The candidacy of YAP1 as a susceptibility gene for PCOS has been highlighted in several studies^15, 108, 109^.
13. **ZBTB16 (Zinc finger and BTB domain containing 16, also known as PLZF): Located at 11q23.1 (Chr 11; 113949232-113949232).** Member of the Krueppel C2H2-type zinc-finger protein family and encodes a zinc finger transcription factor that contains nine Kruppel-type zinc finger domains at the carboxyl terminus. This protein is located in the nucleus and is involved in cell cycle progression. The zinc finger protein has a pro-apoptotic and anti-proliferative activity and has been marked as an androgen-responsive gene with anti-proliferative activity in prostate cancer cells ^23^. PLZF binds to the GATA4 gene regulatory region and activates GATA4 transcription and mediates cardiac hypertrophic signaling from angiotensin II receptor 2^24^. The loss of PLZF has been related to increased proliferation, invasiveness and motility, and resistance to apoptosis in different cancer cell types^110^. PLZF is considered a tumor suppressor gene in various cell types and tissues. Up-regulated during adipocyte differentiation *in vitro*^25^. Involved in control of early stages of spermatogenesis^26^, and critical for endometrial stromal cell decidualization^27^.
14. **ERBB3 (erb-b2 receptor tyrosine kinase 3; also known as HER3): Located at 12q13.** A member of the EGFR family of receptor tyrosine kinases (RefSeq). The ERBB3 gene is a potential susceptibility locus for T1D and has also been associated with PCOS^15, 111^. ERBB4 together with ERBB3-binding protein 1 may modulate the protein cascade that leads to differentiation of ovarian somatic cells. ERBB3 interacts with the YAP protein in the Hippo pathway and is implicated in ovarian cell tumors^112^. The same region also includes RAB5B and a SNP in this region has been associated with response to glycose stimulation^113^.
15. **RAB5B (Member of the RAS oncogene family): Located at 12q13.** Member of the RAS oncogene family. RAB5B is an isoform of RAB5, a member of the small G protein family. Rab5 regulates fusion and motility of early endosomes, and is a marker of the early endosome compartment^114^. Endogenous Rab5B may work in conjunction or in sequence with Rab5A to facilitate the trafficking of EGFR^115^. RAB5b has previously been identified in PCOS in women of Han Chinese and European descent^109^. A variant near this gene has been associated with insulin and glucose levels^113^. It has been suggested that *DENND1A, LHCGR, INSR*, and *RAB5B* form a hierarchical signaling network that can influence androgen synthesis^98^. Proposed to modify PCOS risk through metabolic mechanisms ^76^. RAB5B shows lower expression levels in adipose tissue from PCOS women compared to healthy controls^116^.
16. **KRR1 (KRR1, small subunit (SSU) processome component, homolog (yeast)): Located at 12q21.2 (Chr 12; 75941042-75941042).** Required for 40S ribosome biogenesis. Involved in nucleolar processing of pre-18S ribosomal RNA and ribosome assembly (inferred function based on sequence similarity)^82^. The region also includes the testosterone- and estrogen-sensitive GLIPR1, GLIPR1L1 and GLIPR1L2 genes, which encode proteins involved in male germ cell maturation and sperm-oocyte binding^117–119^. Proposed to modify PCOS risk through metabolic mechanisms^76^.
17. **TOX3 (TOX high mobility group box family member 3): Located at 16q12.1.** This gene regulates Ca2+-dependent neuronal transcription through interaction with the cAMP-response-element-binding protein (CREB)^120^. The protein encoded by this gene contains an HMG-box, indicating that it may be involved in bending and unwinding of DNA and alteration of chromatin structure (RefSeq). The C-terminus of the encoded protein is glutamine-rich due to CAG repeats in the coding sequence. A minor allele of this gene has been implicated in an elevated risk of breast cancer. In normal human tissues, TOX3 is largely expressed in the central nervous system (CNS), in the ileum, and within the brain in the frontal and occipital lobe. TOX3 overexpression induces transcription involving isolated estrogen-responsive elements and estrogen-responsive promoters, and protects neuronal cells from cell death caused by endoplasmic reticulum stress or BAX overexpression^120^. *TOX3* has been highlighted as a potential PCOS susceptibility locus before and there is evidence it may modify the hyperandrogenemic aspects of the syndrome^15, 121^. Proposed to modify PCOS risk through inflammatory mechanisms ^76^.
18. **MAPRE1 (Microtubule-associated protein, RP/EB family, member 1, also known as EB1): Located at 20q11.1-q11.23 (Chr 20; 31420757-31420757).** EB1 interacts with the low-density lipoprotein receptor related protein 1 (LRP1), which controls adipogenesis^28^ and may additionally mediate ovarian angiogenesis and follicle development^29^. EB1 binds to the plus end of microtubules and regulates the dynamics of the microtubule cytoskeleton^82^. It is thought that this protein is involved in suppression of microtubule dynamic instability, regulation of microtubule polymerization and spindle function, and chromosome stability (RefSeq).
19. **ARSD (arylsulfatase D): Located at Xp22.3 (X-chromosome; 2846021-2846021).** ARSD is a member of the sulfatase family and located within a cluster of similar arylsulfatase genes on chromosome X. The encoded proteins are essential for the correct composition of bone and cartilage matrix (RefSeq, GeneNetwork, USCS). This gene has been marked as a prognostic marker in chronic lymphocytic leukemia and has been suggested as a biological mechanism in chroinic lymphocytic leukemia - CLL^122^. The Xp22.3 region also includes the gene for glycogenin 2 (GYG2), which is involved in glycogen biosynthesis and blood glucose homeostasis. It has been shown that glycogen biosynthesis pathways are impaired in PCOS^123, 124^.

#### Supplementary note on gene enrichment analysis

We used MAGENTA (Meta-analysis Gene-set Enrichment of Variant Associations; version 2.4; ^125^ and DEPICT (Data-driven Expression-Prioritized Integration for Complex Traits; release 142 for 1000 Genomes imputed data;^126 127^ methods to specifically prioritize genes, pathways and tissues enriched in the genome-wide results of the PCOS meta-analysis. In brief, MAGENTA assesses the over-representation of genes with low P-values in their locus across manually curated databases. For the MAGENTA analysis, curated gene-sets and Gene Ontology (GO) gene-sets were obtained from the Molecular Signatures Database (MSigDB release v4.0; ^128^). DEPICT prioritizes genes in the associated loci, detects enriched pathways and tissues based on derived data sets based on patterns of co-expression and the expression levels of the genes at associated loci. Further, we performed functional annotation enrichment analysis using GoShifter (Genomic Annotation Shifter; ^129^). Functional annotations used in these analyses were transcription factor binding sites (172 transcription factors)^130^ and chromatin states in different tissues (n=196)^131^. We investigated whether the 14 PCOS-associated susceptibility variants detected in this study and the variants in LD with them (r^2^>0.6) co-localized with specific functional annotations. The results from gene set analysis did not show results that we found particularly trustworthy, and the methods of MAGENTA have been criticized elsewhere so these are only reported in the supplement. No individual pathway appeared to be significant. GoShifter analyses for identification of enriched functional annotations did not reveal any statistically significant finding (all p-value>0.05). DEPICT tissue identification approach reinforced the importance of ovarian morphology, with ovarian follicle, ovum, oocytes, ovary, granulosa cells, fallopian tubes and cumulus cells all showing nominally significant p-values (p-value<0.05). This is alongside the more general enrichment at endocrine cells and adipocytes.

The nominally significant findings for ovaries in the tissue identification analysis suggested the importance of ovarian morphology in PCOS pathogenesis. However, no individual pathway appeared to be significant in gene-set enrichment and gene prioritization analyses. A potential explanation for the lack of significant findings is that these methods are limited by the functional data available for the tissues relevant to PCOS and its related traits, e.g. ovary. In addition, DEPICT and GoShifter analyses were based on the 14 PCOS GWAS meta-analysis susceptibility variants, which may limit the power of these approaches to detect significant enrichments ^129^.

### SUPPLEMENTARY DATA

**Supplementary Tables:**

Supplementary Table 1: *Cohorts contributing polycystic ovary syndrome (PCOS) cases, PCOS phenotypes, laboratory data and controls*.

Supplementary Table 2: Fine-mapping of PCOS risk loci identified in the meta-analysis to narrow candidate causal variants.

Supplementary Table 3: Look-up of previously published PCOS risk variants in Han Chinese cohorts with PCOS GWAS meta-analysis and PCOS related traits (HA, OD, PCOM, T, FSH, LH and ovarian volume).

Supplementary Table 4: Look-up of PCOS GWAS meta-analysis susceptibility variants with PCOS related traits (HA, OD, PCOM, T, FSH, LH and ovarian volume).

**Supplementary Figures:**

**Supplementary Figure 1.**
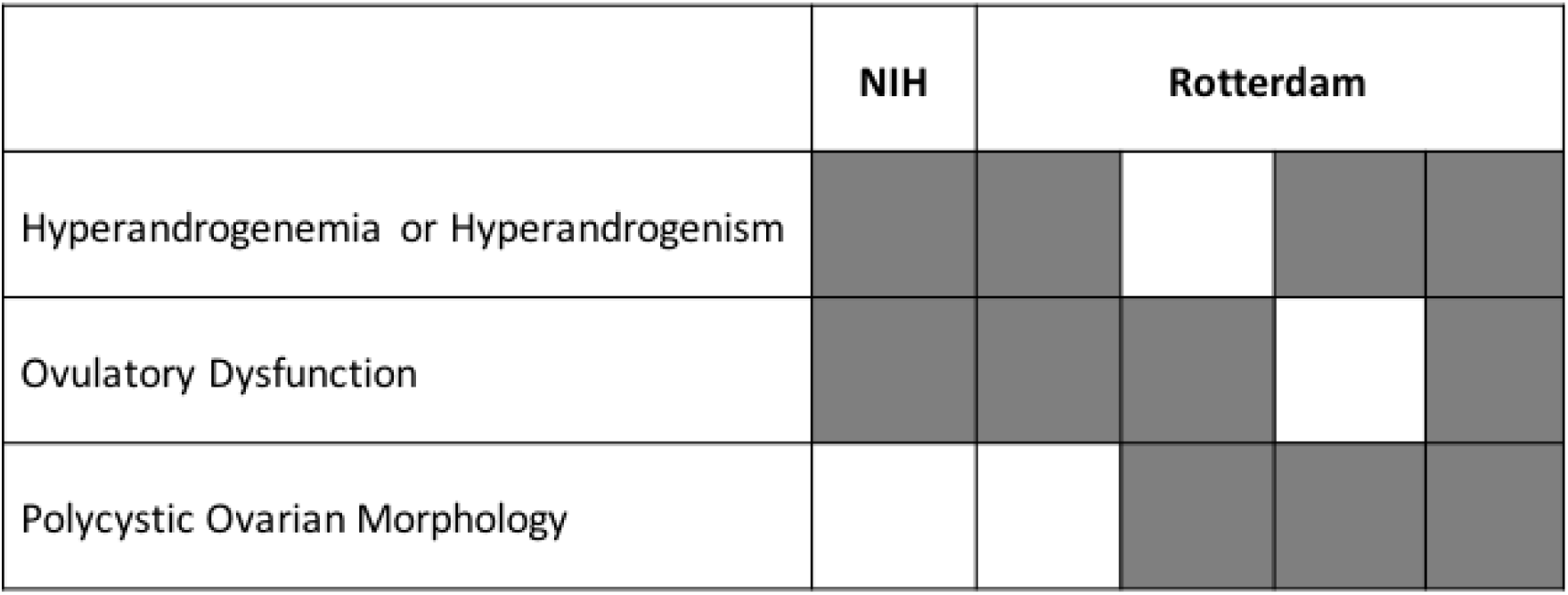
Diagnostic criteria of PCOS. Columns represent the diagnostic phenotypes that result from different diagnostic criteria. Grey squares indicate required traits for diagnosis within each diagnostic phenotype.

**Supplementary Figure 2.**
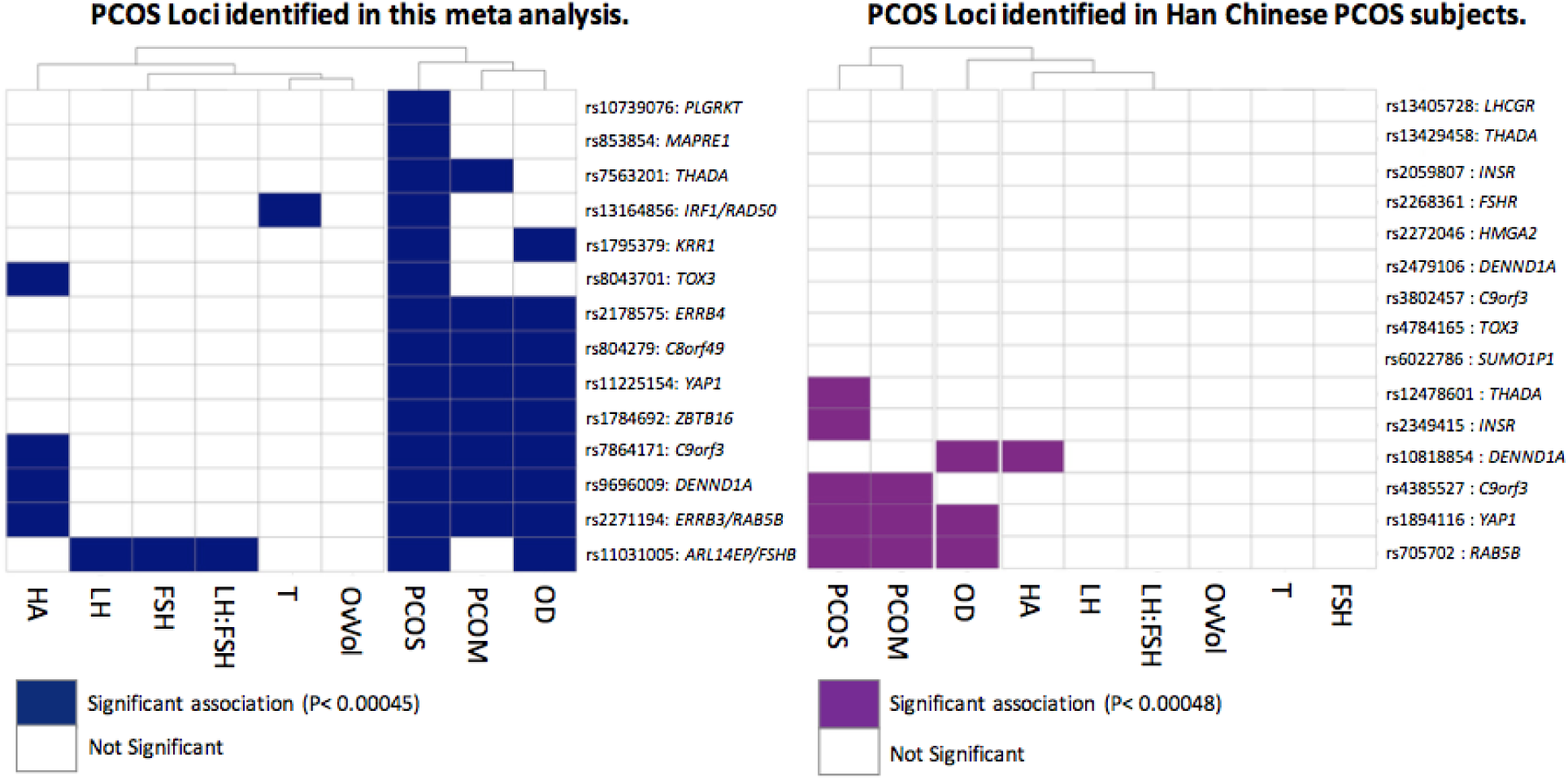
Cluster plots showing relationships between PCOS loci and related traits. Loci significantly associated with PCOS in our meta analysis (shown on left, blue) or in the previously reported meta analyses of Chinese PCOS subjects in the analysis of related traits in our own meta (shown on right, purple). Clustering by column (phenotype/trait) demonstrates the large proportion of PCOS loci that are also significantly associated with ovulatory dysfunction (OD) and polycystic ovarian morphology (PCOM).

**Supplementary Figure 3:**
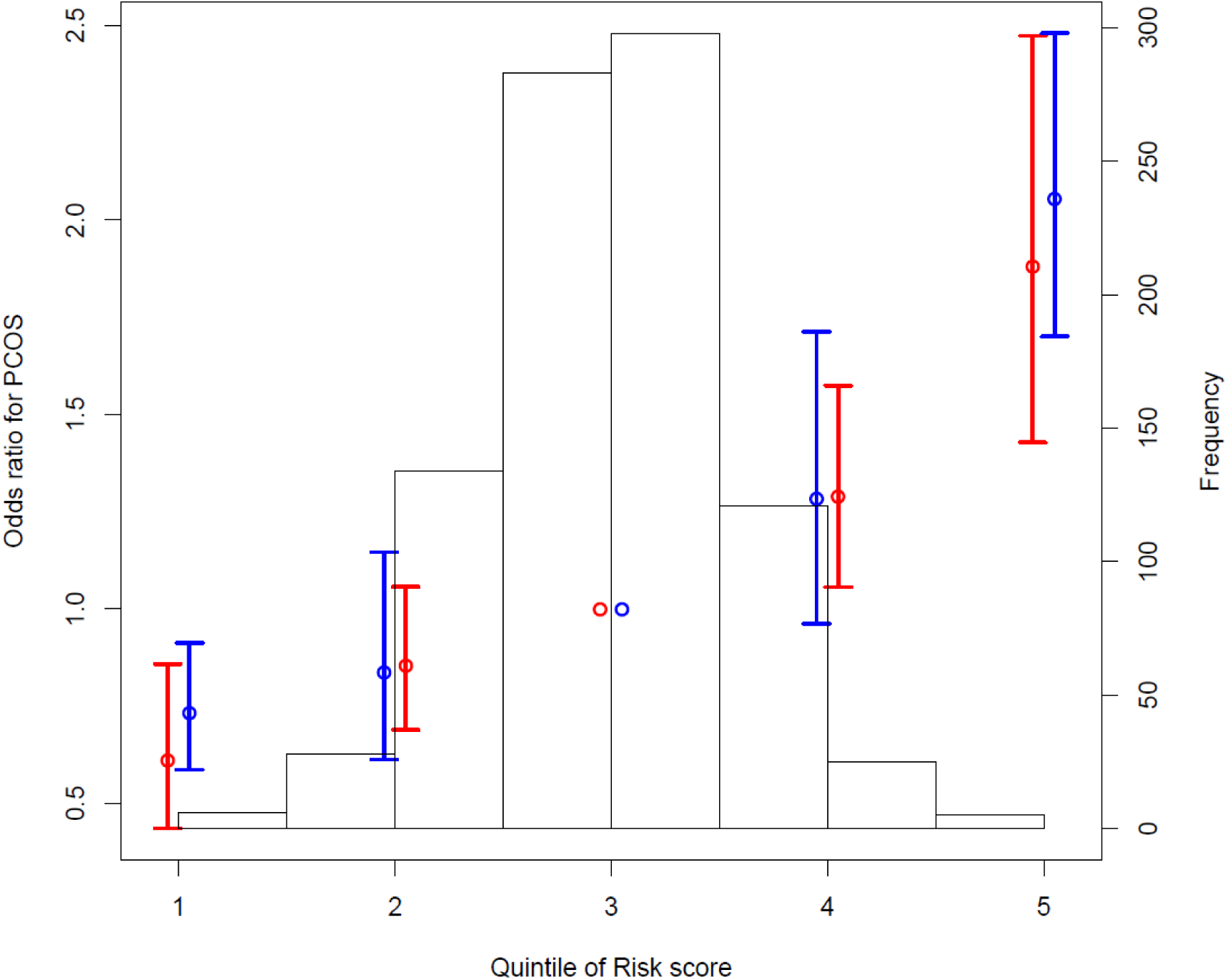
Weighted genetic risk score. The odds for having PCOS by Rotterdam (blue) or NIH (red) diagnostic criteria based on genetic risk score from across the identified genome-wide significant loci. The group with the average number of risk alleles was used as the reference group.

